# Dynamics of influenza transmission in vampire bats revealed by longitudinal monitoring and a large-scale anthropogenic perturbation

**DOI:** 10.1101/2024.07.26.605290

**Authors:** Megan E Griffiths, Alice Broos, Juan Morales, I-Ting Tu, Laura Bergner, Abdelkader Behdenna, William Valderrama, Carlos Tello, Jorge E Carrera, Sergio Recuenco, Daniel G Streicker, Mafalda Viana

**Author notes:** Joint senior authors (Daniel G Streicker, Mafalda Viana).

## Abstract

Interrupting pathogen transmission between species is a priority strategy to mitigate zoonotic threats. However, avoiding counterproductive interventions requires knowing animal reservoirs of infection and the dynamics of transmission within them, neither of which are easily ascertained from the cross-sectional surveys which currently dominate investigations into newly discovered viruses. We used biobanked sera and metagenomic data to reconstruct the transmission of recently discovered bat-associated influenza virus (BIV) over 12 years in three zones of Peru. Mechanistic models fit under a Bayesian framework, which enabled joint inference from serological and molecular data, showed that common vampire bats maintain BIV independently of the currently assumed fruit bat reservoir through immune waning and seasonal transmission pulses. A large-scale vampire bat cull targeting rabies incidentally halved BIV transmission, confirming vampire bats as maintenance hosts. Our results show how combining field studies, perturbation responses and multi-data type models can elucidate pathogen dynamics in nature and reveal pathogen-dependent effects of interventions.

## Introduction

Pathogen transmission between species (‘spillover’) is responsible for emerging infectious diseases affecting wildlife conservation, agriculture, and human health. Understanding the ecological and biological mechanisms that enable pathogen maintenance within the reservoir hosts is a prerequisite to anticipating emergence risk and developing interventions to prevent spillover (1). However, identification of reservoir hosts is notoriously difficult since not all susceptible hosts are important for transmission (2,3). For example, SARS-CoV-2 nucleic acids or antibodies have been detected in more than 50 species, but evidence of onward transmission under natural conditions is limited to just 3 species aside from the putative bat hosts (humans, mink, and white-tailed deer) (4–6). Reservoir identification requires long term data from multiple populations and, ideally, evidence that interventions in a hypothesized reservoir predictably alter pathogen dynamics, both of which are rarely available in wildlife systems (7,8). With the reservoir identified, the within and between host biology of pathogens can then be attained using data-driven mathematical models (9,10). To date, such models have been parameterized from population-level serological or molecular pathogen detection data, sometimes constraining parameters with data from other sources (e.g. experimental infections) (11–13). However, the general sparsity of long-term data from wildlife, particularly from replicate populations, means that models often fail to confidently distinguish between different hypothesized models of transmission. Efforts to formally integrate serological and molecular data or to harmonize data collected at the population level (i.e., prevalence or seroprevalence) and individual level (i.e., seroconversion or infection histories of known individuals) remain rare, but have been suggested to improve model performance and ecological understanding (2).

Influenza A viruses (IAV; family Orthomyxoviridae) have caused four major human pandemics, either as a result of spillover from wild bird reservoirs or via intermediate hosts such as domestic pigs (14–16). The discovery of bat-associated influenza A viruses (BIVs, subtypes H17N10 and H18N11) in 2012-2013 revealed neotropical bats as a potential source for influenza emergence, catalysing an explosion of virological research (17–19).

Modern capacity to sequence viral genomes and rescue infectious clones demonstrated, for example, that cell entry is mediated by MHC-II receptors instead of the canonical human or avian sialic acid linkage receptors (20,21), that BIVs can use human MHC for cell entry and replicate in canine cell lines, and that they transmit efficiently among captive bats, but replicate poorly in mice and ferrets (22). In contrast, knowledge of the putative reservoir hosts, and of the ecology and epidemiology of BIVs, is largely speculative. Current understanding is derived from three field surveys which detected BIV nucleic acid in rectal swabs from a total of 6 individuals of three species (*Sturnira lilium* in Guatemala, *Artibeus planirostris* in Peru, and *Artibeus lituratus* in Brazil) and a single cross-sectional survey which detected BIV antibodies in 13 of 18 species (17,18,23). As such, which bat species are important for BIV transmission and how the incidence of infection may vary seasonally or through space and time remain unknown.

Here, we characterized the transmission dynamics of BIVs in populations of a putative reservoir, the common vampire bat (*Desmodus rotundus*, family Phyllostomidae). In Peru, BIV antibodies were previously detected in approximately 40% of vampire bats, though none of the 18 bats were detectably shedding virus at the time of capture (17). The obligately blood feeding nature of vampire bats means contacts with humans, livestock and wildlife during blood feeding are frequent (24,25), opening the possibility of direct viral transmission to other species via biting (akin to rabies virus) or through exposure to urine or faeces during feeding. If vampire bats transmit BIV competently, such frequent opportunities for spillover may provide a foothold for host adaptation and onwards transmission in non-bat species (22,26–29). Vampire bats also span the known geographic range of BIVs and occur within both undisturbed and anthropogenically transformed ecosystems. Uniquely among bats, vampire bats are culled using species-specific, spreadable anticoagulant poisons with the intention to reduce the incidence of bat bites on humans and livestock and lethal rabies virus spillover. However, evidence to date suggests these culls fail to reduce the rabies incidence and may exacerbate spatial spread owing to the higher sensitivity of rabies transmission to bat movement than to bat density (11,30). How BIVs and other pathogens respond to vampire bat culls is unknown.

To evaluate the potential for vampire bats to transmit BIV and to interrogate the within-host and ecological mechanisms of long-term viral maintenance, we fit compartmental models of BIV transmission to spatially replicated field data from *D. rotundus* spanning from 2007-2018 from three zones of Peru (Figure 1). First, we contrast competing models of BIV biology focusing on whether infection leads to lifelong or transient immunity. Because our dataset included several hundred bats that were re-captured and re-sampled at irregular intervals and cross-sectional metagenomic sequencing (31), we use a Bayesian modelling framework to directly integrate these individual-level and molecular surveillance data, showing that this joint approach improves parameter estimation over models that exclusively use population-level serological data. Next, we ask whether the seasonal and inter-annual dynamics of infection varied across three zones of Peru with distinct environmental conditions and assess how this variation impacts the expected frequency of infectious bats. Finally, using data from a geographically expansive bat cull carried out over 2 years (32), we explore whether a real world intervention reduced BIV transmission, as might be expected if vampire bats are competent maintenance hosts of H18 influenza (8).

**Figure 1.**
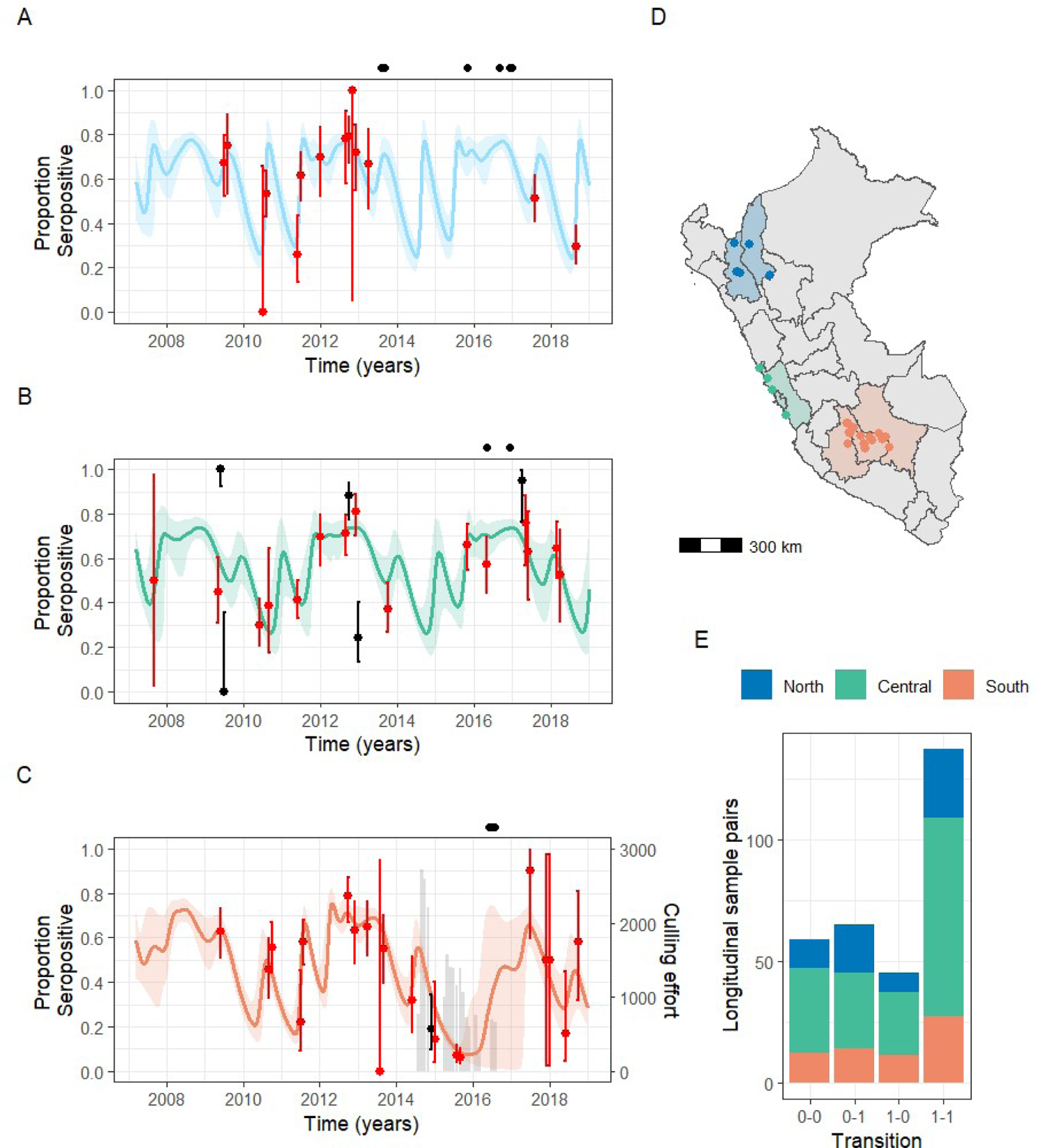
Observed and model-predicted seroprevalence of influenza virus across three zones of Peru. In (A)-(C), solid lines show the mean predicted trajectory for the recovered class in an SEIR model. Shaded regions show the 95% credible intervals (CIs) for the (A) North, (B) Central and (C) South zones. Points show monthly population-level seroprevalence data with binomial confidence intervals, with observations that fell outside the 95% CI of the posterior cumulative distribution shown in black. The grey bars in panel (C) show monthly applications of vampiricide during a culling campaign in the South. Black points above each time series show the collection dates for samples subjected to metagenomic sequencing. (D) Colours show the administrative regions of Peru which were grouped for this study, and the points show the locations of sampled bat colonies. (E) A summary of change in serostatus in the paired samples from longitudinally sampled bats where 0 = seronegative and 1 = seropositive.

## Results

A total of 2551 serum samples were collected from 2460 vampire bats between 2007 and 2018, from 30 bat roosts in 3 ecological zones of Peru (Figure 1). Of these, 16 roosts contained only *D. rotundus* and 14 also contained other bat species including multiple *Artibeus* species in which BIV H18 RNA has been previously detected (Table S1 (17)). Males and females were sampled relatively evenly (52.4% and 47.6% respectively), and most bats were adults (88.5%). Using an ELISA targeting H18 BIV, approximately half of the samples were seropositive (52.3%) with average seropositivity higher in the ‘Central’ zone (59.4%; N=958) and the ‘North’ zone (55.9%; N=614) compared to the ‘South’ zone (36.3%; N=979). Seroprevalence was high in vampire bats in the presence (64.3%, binomial CIs: 61.1-67.5%) and absence 51.6% (49.0-54.3%) of co-roosting *Artibeus* species.

A total of 269 individually marked vampire bats were recaptured and sampled at least twice during the study period (range: 2-4 captures, time between sampling: 3-96 months). Of 306 sample pairs, 44.8% were seropositive at both timepoints, 21.2% seroconverted (0–1) and 14.7% showed complete antibody waning (1–0) (Figure 1E). Of the bats sampled >2 times (N=37), 5 were seropositive after previous antibody waning (1-0-1), suggesting reinfection or re-exposure to BIV boosted antibody production (Figure S1). As previous field and experimental challenge studies detected BIV RNA in bat faeces, we mined existing metagenomic sequencing data from the same vampire bat populations for the presence of viral RNA. Reference alignment of sequencing reads to a published H18N11 genome (CY125949 (17)) detected no BIV RNA in 18 sequencing pools derived from 172 bat faecal samples (Figure 1).

### Comparing models of BIV immunity suggests immune waning and re-infection

Antibody loss in formerly seropositive individuals (Figure 1E) could renew susceptibility to future BIV exposures or could indicate continued protection via non-antibody (e.g., cellular) mediated immunological mechanisms or anamnestic responses (33). To distinguish these hypotheses of waning and lifelong immunity, we fit competing epidemiological models which assumed either that bats return to a susceptible (S) class after a period of immunity in a recovered class (R) (‘waning immunity’) or retain lifelong immunity despite loss of detectable antibodies (‘lifelong immunity’) (Figure S2). In both models, bats are born susceptible (S), become exposed (E) to BIV at transmission rate β, become infected (I) after an incubation period (1/ δ) and clear BIV at rate 1/α to enter the recovered (R) compartment (see Methods). In model 1, R bats (assumed seropositive) return to the S at an antibody waning rate of 1/γ. In model 2, seropositive bats (R^+^) lose detectable antibodies at rate 1/γ but remain immune (R^-^). These bats can become seropositive again due to re-exposure to BIV upon contact with an infected bat at rate β (i.e., boosting, Figure S2). In both models, the transmission rate β includes annual (e.g., seasonal births or dispersal) and multi-year periodicity (e.g., bat demography, climatic events). Parameters α, δ, and γ were assumed to be constant across zones as intrinsic virus replication cycle properties, and thus estimated jointly, with γ estimated through an occupancy model simultaneously fitted to the serological data from recaptured bats (Figure 1E). Transmission parameters (Equation 11) were estimated separately for the three datasets corresponding to the sampled zones of Peru. In the ‘South’ zone a geographically extensive vampire bat cull took place during our study period which dramatically reduced vampire bat population sizes, and visually, appeared to be associated with fluctuations in H18 seroprevalence (32) (Figure 1C). We therefore included a culling term in the South model and evaluate the effects of this decision in later analyses.

Our model comparisons supported immune waning over lifelong immunity. The waning immunity model had a lower Deviance Information Criterion (ΔDIC = 86.32) and Leave-one-out Information Criterion (ΔLOOIC = 146.10), converged better (maximum Gelman-Rubin diagnostic of 1.02 vs 2.76 when assuming lifelong immunity) and fitted better to the BIV seroprevalence in each zone, with ∼90% of data points falling inside the 95% CI of the cumulative distribution of the posterior (Figure 1A-C; Table 1) compared with the marginal reduction to ∼85% in when assuming lifelong immunity (Figure S3). Hereafter, unless otherwise noted, results use the better supported model of immune waning. The absence of lifelong immunity points to re-infection as a potentially important mechanism for long-term influenza maintenance in bats.

**Table 1.** Final model full parameter priors and posteriors. For all parameters estimated separately for each zone, results are presented in the order North, South, Central.

| Parameter | Definition | Estimated | Distribution | Prior | Reference | Posterior mean | Posterior SD | Effective sample size | Rhat |
| --- | --- | --- | --- | --- | --- | --- | --- | --- | --- |
| b and d | Lifespan (years) | No | - | 3.5 | (44) | - | - | - | - |
| $\delta$ | Incubation period (days) | Yes | Gamma | Mu=3<br>Var=1.5 | (22,41) | 1.99 | 0.70 | 3355 | 1.00 |
| $\alpha$ | Infectious period (days) | Yes | Gamma | Mu=7<br>Var=14 | (22,41) | 6.12 | 0.72 | 1315 | 1.00 |
| $\gamma$ | Immune period (days) | Yes | Gamma | Mu=365<br>Var=10,000 | NA | 232.43 | 30.21 | 596 | 1.02 |
| beta0 | Baseline in absence of periodicity | Yes | Gamma | Mu=8<br>Var=8 | NA | 5.00 | 1.07 | 4000 | 1.00 |
|  |  |  |  |  |  | 7.97 | 1.32 | 2803 | 1.00 |
|  |  |  |  |  |  | 5.83 | 1.17 | 610 | 1.01 |
| beta1 | Impact of short-term periodicity | Yes | Beta | Mu=0.4<br>Var=0.04 | NA | 0.55 | 0.16 | 3716 | 1.00 |
|  |  |  |  |  |  | 0.31 | 0.09 | 2653 | 1.00 |
|  |  |  |  |  |  | 0.31 | 0.12 | 584 | 1.00 |
| beta2 | Impact of long-term periodicity | Yes | Beta | Mu=0.5<br>Var=0.04 | NA | 0.72 | 0.13 | 3293 | 1.00 |
|  |  |  |  |  |  | 0.69 | 0.08 | 4000 | 1.00 |
|  |  |  |  |  |  | 0.72 | 0.10 | 1397 | 1.00 |
| period 1 | Short-term period (days) | No | - | 365 | NA | - | - |  | - |
| period 2 | Long-term period (days) | Yes | Uniform | Min=365*3<br>Max=365*5 | NA | 1424.35 | 28.69 | 4000 | 1.00 |
|  |  |  |  |  |  | 1579.80 | 50.15 | 3211 | 1.00 |
|  |  |  |  |  |  | 1459.16 | 54.12 | 1143 | 1.00 |
| x1 | Timing of short-term periodicity | Yes | Uniform | Min=-1<br>Max=1 | NA | 0.35 | 0.11 | 2900 | 1.00 |
|  |  |  |  |  |  | 0.81 | 0.11 | 1460 | 1.01 |
|  |  |  |  |  |  | -0.24 | 0.13 | 4000 | 1.00 |
| x2 | Timing of long-term periodicity | Yes | Uniform | Min=0<br>Max=2 | NA | 0.74 | 0.08 | 1682 | 1.00 |
|  |  |  |  |  |  | 0.84 | 0.09 | 1200 | 1.01 |
|  |  |  |  |  |  | 0.74 | 0.12 | 870 | 1.00 |
| csf | Culling scaling factor | Yes | Gamma | Mu=5E-3<br>Var=1.25E-5 | NA | 0.0023 | 8.30E-4 | 4000 | 1.00 |

### Serological data from multi-captured bats refines the within-host dynamics of BIV

We assessed the relative value of adding individual-level seroconversion history data from recaptured individuals and metagenomic data to the “core” population-level serological data, for identifying the parameters that govern the within-host dynamics of BIV. We first compared the theoretical performance of our model integrating these three types of data with that using the core data only, by simulating data and comparing the estimated parameters with the true input values. As expected, full data integration improved model performance, with the individual-level serology data providing the most additional information (Figure S4).

Using the full dataset, we estimated an incubation period of 2.0 days (95% CI: 1.1-3.7), and an infectious period of 6.1 days (CI: 4.9-7.8), with minimal difference when estimated using only the population-level serology data (Figure 2). Our model estimated the immune period γ at 232.4 days (CI: 187.7-310.8). Individual-level serological data were highly influential for estimates of γ, with model fitting using only the population-level data shortening the immune period by 49.0 days (183.4 days, CI: 147.5-224.2; Figure 2 grey lines, Figure S5). Given that our earlier simulations showed that γ was the parameter that was most significantly and consistently improved by data integration (Figure 2C; Figure S4), we argue that the higher estimate from the full dataset should be closer to the true value. This demonstrates that for the first full epidemiological characterisation of BIV in the field, combining different sample types and individual and population scales of observation can compensate for a lack of long-term experimental data in captive bats, insufficient to observe antibody waning.

**Figure 2.**
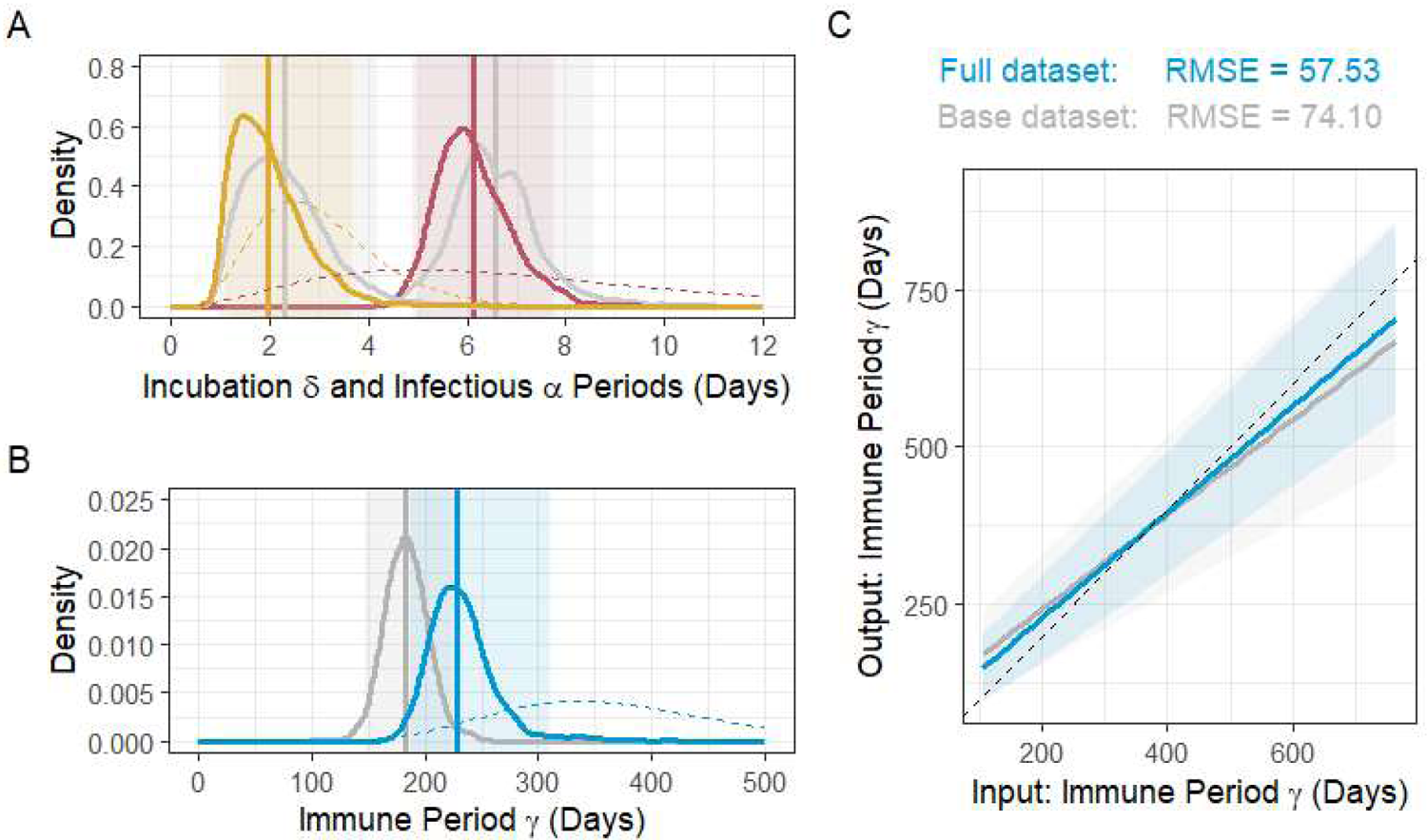
Inference from multiple data types refines the within host biology of BIV. Prior distributions (dashed) and posterior distributions for (A) δ (incubation period; yellow) and α (infectious period; red), and (B) γ (immune period; blue) produced by model fitting to the full dataset including population level serology, individual-level serology from recaptures, and metagenomic sequencing. Grey lines show the posterior distribution for each parameter when fitted to the base dataset of only the population-level serology data. The mean of each posterior distribution is shown by a solid vertical line, and 95% CIs are shown by the shaded areas. Full comparison of data inclusion can be found in Figure S5. (C) Model fitting to simulated data comparing the full dataset and the base dataset. The x-axis of each graph shows the input parameter value, and the y-axis shows the mean of the posterior distribution when model fitting to test data. The shaded areas show 95% credible intervals.

### Population dynamics of BIV: Regionally distinct peaks in infection prevalence

Our model evaluation and estimation of within-host parameters support a short immune period as important for BIV transmission and maintenance. However, assessing the potential for spillover requires that we understand how transmission and infection prevalence vary at finer scales; therefore, we next explored the spatial and temporal variation in BIV dynamics. Both annual and inter-annual periodicity parameters (β_1_ and β_2,_ respectively) had posterior means and lower credible bounds >0, supporting seasonality and multi-year cycles of BIV transmission within each zone of Peru (Figure 3A; Figure S7). Inter-annual cycles had an approximately 4-year period and were largely synchronous between zones. In contrast, seasonality was asynchronous across zones (Figure 3C). Infection within the South zone peaked the earliest from March-May each year, with the exception of 2015 in which transmission remained low. In the North zone, transmission peaked from mid-May-early July, with some overlap between this peak time and that in the South. The infection peak in the Central zone tended to occur later in the year from September-October, with no overlap with the other zones. Taken together, these results imply that broad scale drivers of BIV transmission may harmonize inter-annual dynamics across zones, but that ecological differences within zones shape local transmission dynamics.

**Figure 3.**
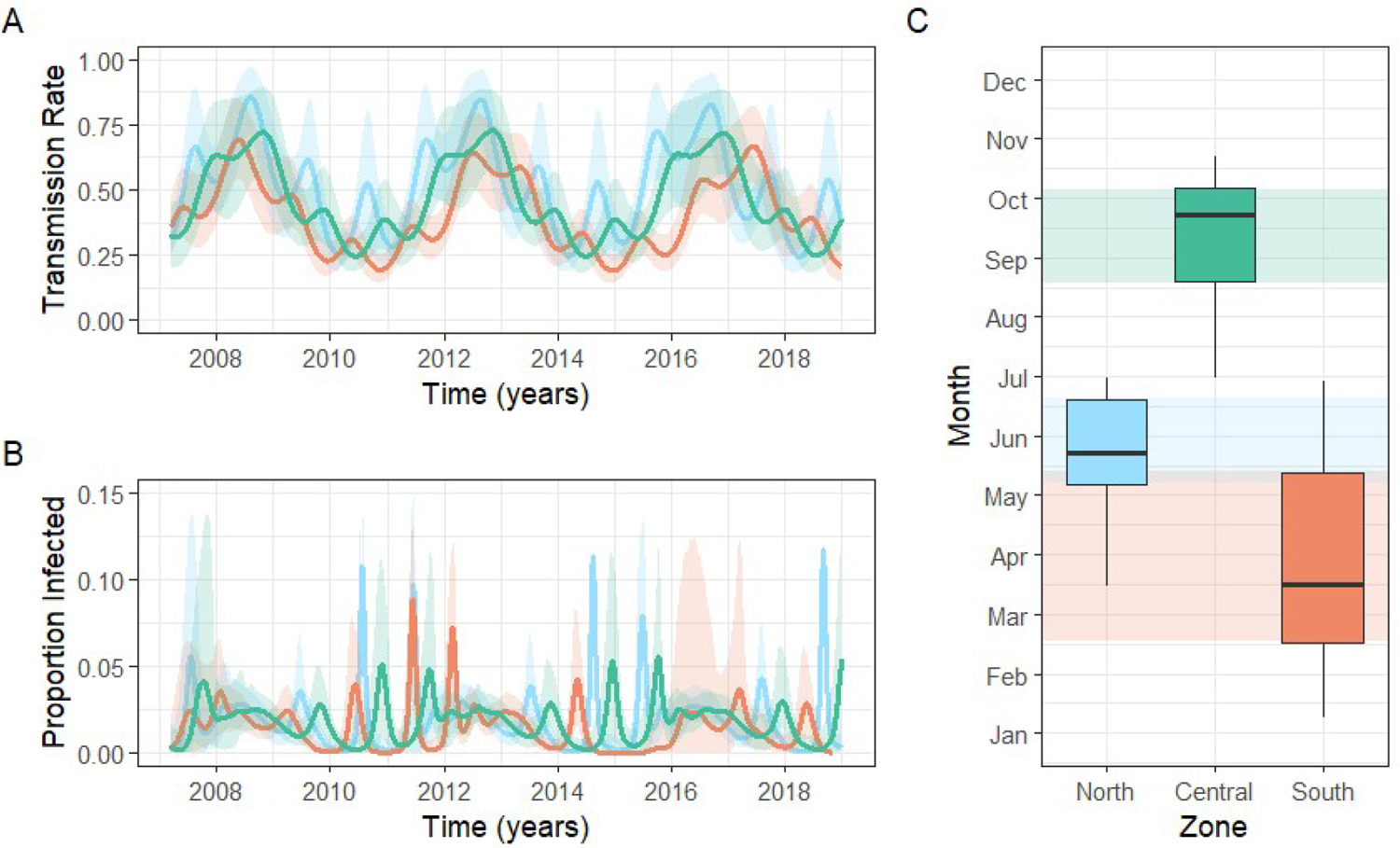
Regionally synchronised long-term dynamics with distinct seasonal peaks of BIV transmission and infection. Trajectories for (A) the time varying rate of transmission, beta, and (B) the proportion of the population actively infected by BIV predicted by model fitting for each of the three zones of Peru. Solid lines show the mean trajectory, and shaded areas show the 95% CIs. (C) Boxplot showing the predicted seasonal peak of transmission by zone.

We next used our models to estimate the incidence of active BIV infection in vampire bats through time. The mean predicted infection prevalence across Peru was <2%, consistent with the infrequent detection of BIV by our metagenomic sequencing and by RT-PCR in previous studies (17,18,23). Spatial variation in the mean percentage of infected bats was minimal (North, 1.9%; Central, 1.8%; South, 1.4%); however, dramatic peaks in transmission varied by zone. For example, we estimated that 11.5% of the vampire bat population was infectious and shedding BIV in the North mid-2018 (Figure 3B). In contrast, peaks of infection never exceeded 6% of the population in the Central zone. From an applied standpoint, these spatiotemporally explicit predictions of BIV infection prevalence should correspond to exposure risks in non-bat species and might inform priority sampling of locations and times when virus detection by molecular methods or isolation would be fruitful.

Dynamical evidence of sustained transmission in a single species model suggested the potential for vampire bats to maintain BIV transmission over multi-annual cycles independently of other bat species, including the suspected *Artibeus* reservoir. However, these results derived from models which do not easily allow for viral extinction. To assess the potential for vampire bats to indefinitely maintain BIV, we therefore ran stochastic simulations of BIV transmission using the posterior mean estimates of each parameter and measured the proportion of simulations which led to viral extinction (no exposed or infected individuals present) at the end of the 12-year time period. Crucially, our models predict that BIV should persist in vampire bat population sizes that can be reasonably expected within the zones analysed here (e.g., <15,000 bats are required for 95% BIV survival for 12 years in the Central zone), supporting their potential to maintain BIV independent of other bat species (Figure S6).

### Impacts of bat culls support vampire bats as a maintenance host of BIV

Our model supported the possibility that vampire bats maintain BIV independently of other bat species. Another important, but rarely attained, line of evidence for identifying reservoir hosts is whether landscape scale interventions impact transmission within the suspected reservoir (8). We opportunistically assessed whether a geographically expansive vampire bat cull, which was intended to suppress rabies transmission in the South zone of Peru, affected BIV transmission ((32), Figure 1C). Importantly, this cull used an anticoagulant poison (vampiricide) which was applied topically to captured vampire bats and spread between them by allo-grooming, making impacts on non-vampire bat species unlikely (34). We investigated the impact of culling by estimating an additional parameter ‘*csf’* within our Bayesian epidemiological model. *csf* works as a scaling factor for the impact of the culling, where *csf*=0 would mean no effect of culling on BIV transmission (as would be expected if BIV exposures in vampire bats were density independent or driven by non-vampire bat species unaffected by culls), and higher values correspond to negative effects of culling on BIV transmission. This factor is applied to all classes in the model, i.e., culling proportionally reduces the population size in each class. The model predicted a substantial reduction of

BIV transmission due to vampire bat culling (posterior mean *csf* = 2.3E-3; 95% CI: 0.99E-3 – 4.1E-3; Figure 4A). This suggests that the interruption of long-term periodicity in the South relative to other zones and the decreased magnitude of the expected the annual peak in Infecteds in 2015 and 2016 (Figure 4B), may have been attributable to bat culls. Comparing the estimated proportion of BIV infected bats and the predicted trajectory in the hypothetical absence of culling showed that culling reduced the incidence of BIV infection by 54.5% (Figure 4B). This reduction supports *D. rotundus* as a reservoir host of BIV, with the observed seroprevalence driven by intra-species transmission events as supposed to frequent spillover from co-roosting bat species.

**Figure 4.**
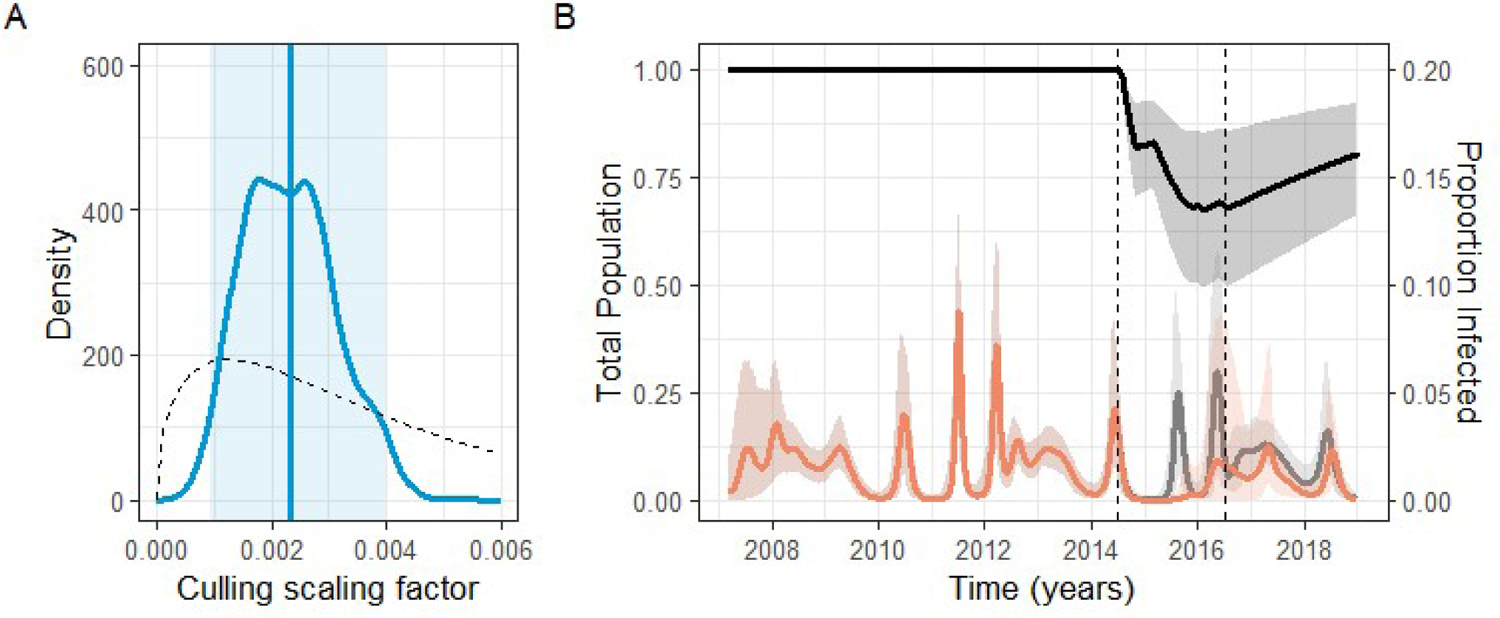
Culling vampire bats reduces BIV transmission. (A) Posterior (solid blue line) and prior (dashed line) distributions of the culling scaling factor (csf) evaluating the impact of culling on BIV transmission. The mean of the posterior is shown by the vertical blue line, and the 95% credible interval by the shaded area. (B) The predicted trajectory for the total vampire bat population in the South zone, as a proportion of the starting population. The mean predicted trajectory is shown by the solid black line, and the 95% credible interval by the shaded area. The predicted trajectory for the Infecteds in the presence (orange line) or absence (grey line) of culling is shown as a proportion of the population. Vertical dashed lines show the start and end dates of the culling campaign.

Our BIV transmission model also estimated the magnitude of vampire bat population reduction during the culling program, which was unknown because poison was applied to 20,957 common vampire bats (35), but neither the total vampire bat population size nor the extent of bat-to-bat spread of the poison was monitored. This second point is especially confounding, as estimates of the spreadability of vampiricide vary greatly, from 1.5-15x the number of bats treated with poison (36,37). The mean population trajectory estimated by the model shows a maximum decrease in the total population size of 32.4% (CI: 14.8%-50.5%; Figure 4B) due to culling, with the lowest population predicted for December of 2015, towards the end of the culling period. Despite this reduction, culling was insufficient to eliminate BIV, with seroprevalence and predicted incidence rapidly returning to pre-culling levels after the cessation of culls.

## Discussion

Cross-sectional studies into the circulation of pathogens of zoonotic concern in their natural hosts typically fail to resolve which infected hosts act as reservoirs or how spillover risk varies in space or time. For neotropical bat-associated influenza viruses, field surveys have detected, albeit rarely, evidence of viral shedding in three frugivorous bat species, leading to the presumption that these species are viral reservoirs. However serological evidence indicated susceptibility in a variety of other bat species (17,18,23). By analysing spatially or spatiotemporally replicated serological and molecular data on bat influenza infection and responses to a large-scale anthropogenic perturbation, we identify common vampire bats as a likely reservoir of BIV and show that long-term viral maintenance in this species is driven by immune waning, seasonal transmission pulses, and synchronized inter-annual cycles.

Several lines of evidence support vampire bats act as a reservoir host of BIV. First, seroprevalence in vampire bats was consistently high across the 12-year period studied here, including within colonies containing *D. rotundus* only (16/30), where seropositivity is less plausibly explained by exposures from alternative bat hosts. This included vampire bats on the Pacific coast of South America, where none of the bat species occur from which viral RNA was previously detected. Second, our stochastic simulations (Figure S6) suggested that extinction of BIV would be unlikely within the range of vampire bat population sizes expected within our sampled zones (38). Third, vampire-bat specific culling was linked to a substantial decrease in BIV seropositivity, indicating a relationship between vampire bat density and BIV transmission which would not be expected if vampire bats were incidental hosts (39).

Our results therefore suggest that vampire bats are sufficient for long term BIV maintenance. However, it remains unclear whether vampire bats form a single species reservoir or form part of a more complex reservoir community containing multiple competent host species, akin to the involvement of multiple bird species in avian influenza A viruses (40). Anecdotally, vampire bats are found throughout the known geographic range of BIV, indicating this species as a common denominator for BIV presence. The lack of detection of viral shedding in wild vampire bats may reflect a combination of the time varying infection rates demonstrated here, uncertainties in virus shedding pathways, small sample sizes tested with targeted assays (i.e., RT-PCR, N<100 vampire bats tested across three prior studies) and the lower sensitivity of untargeted metagenomic sequencing used here. To date, experimental infections with BIV have used frugivorous bat species which are related -but not identical - to the species which have been RT-PCR positive in nature (22,41). Our results call for *in vivo* assessments of the infection dynamics and transmission competency of vampire bats against BIV and highlight a potential risk of primacy biases, wherein the first host species with confirmed infections are presumed to play disproportionate roles in transmission, narrowing subsequent scientific focus on these species.

Vampire bats appear to maintain BIV through seasonal transmission and re-infection enabled by immune waning. Both the duration of the antibody response to BIV and the relationship between this response and functional immunity, were key unknowns from prior experimental infection studies (22,41–43). Our findings based on seroprevalence and seroconversions in recaptured individuals indicates that the immune period lasts under 8 months. This implies that immunity wanes sufficiently rapidly to replenish the pool of susceptible bats annually (44), such that individuals may undergo multiple re-infections during their lifetimes, consistent with our longitudinal re-capture data (Figure S1). We also observed seasonal peaks in BIV transmission which were asynchronized among the three eco-zones of Peru that we studied. Seasonal births affect the dynamics of variety of wildlife diseases by providing a pulse of susceptible hosts and have been reported in vampire bats (45–48). Although spatial variation in seasonal births has not been explored, both the magnitude and timing of birth seasonality may differ considering the major environmental differences in our study areas, which spanned coastal deserts, inter-Andean valleys, and tropical rainforests with different seasonal patterns in temperature and precipitation. In contrast to seasonality, inter-annual periodicity was synchronized among zones, suggesting epidemiological connectivity. This putative regional linkage is unlikely to be mediated by vampire bat movement since genetic studies of *D. rotundus* in Peru suggests this species rarely travels long distances at the rates that would be required to produce the observed patterns (49,50). Alternatively, long distance migrants (e.g., *Nyctinomops*, and *Lasiurus spp.*) may facilitate virus spread at regional scales, although BIV surveys in these species are lacking (51). For avian influenza virus, phylogenetic studies of viruses from different bird species have demonstrated the role of migratory birds spreading avian influenza to resident, non-migratory birds (52,53). Similar studies of viral genetic diversity in vampire bats and co-roosting bat species would be insightful to confirm the connectivity indicated by our models and reveal the potential host species linked to spatial spread.

Regardless of the underlying ecological drivers, the regular peaks in BIV prevalence predicted by our models provide opportunities for cross-species exposures, especially considering the nightly contacts between vampire bats and their prey, which include IAV susceptible species such as pigs, cows, poultry, marine mammals and humans (54–56). That no overt evidence of BIV spillover to vampire bat prey has been observed may imply that molecular barriers preclude transmission. Indeed, whilst the host tropism of BIV is conceivably broad due to the ubiquity of MHC-II, viral particles produced via reverse genetics failed productively infect human cell lines (57–59). Alternatively, cross-species infections may have gone undetected if not associated with clinical disease or because of inadequate surveillance. Our models can be used to direct active surveillance of humans and animals including livestock towards predicted high-risk periods to more efficiently determine whether spillovers from bats have occurred undetected.

Field studies of wildlife diseases often generate a mixture of serological and molecular data which are collected cross-sectionally or at irregular intervals due to financial constraints and the availability of diagnostic tools. Classically, these datasets have been analysed independently, despite containing potentially complementary epidemiological information.

Our Bayesian framework that integrates three data types (population level seroprevalence, seroconversion histories in individual bats, and snapshot metagenomic sequencing) enabled triangulation of key epidemiological parameters. Importantly, it allowed a flow of information between scales of observation, with the individual-level model taking data from the population-level model, with both models informing parameter estimation. Specifically, incorporating seroconversion histories extended the duration of immunity, a parameter was unknown from experimental infections because of the necessarily short duration of those studies (Figures S4 and S5) (22,41). The addition of metagenomic data had less dramatic effects on epidemiological dynamics or parameter estimation, presumably because the absence of BIV shedding observed was consistent with the signal in the more abundant serological data in this system. Nevertheless, our approach provides a general framework to incorporate virus detection data from metagenomics, PCR, or virus isolation into mechanistic modelling, even when sparse and collected at different time scales from more commonly modelled serological data.

Recent events (e.g., COVID-19 pandemic and H5N1 outbreaks) have increased the appreciation for the importance of preventing initial instances of virus transmission from reservoirs to other species. Proposed approaches advocate both non-specific interventions such as conservation or restoration of wildlife habitats in the hopes that improved wildlife health will reduce spillover risk, and targeted approaches using scalable wildlife vaccination to reduce the incidence of specific pathogens in wildlife reservoirs (60,61). Our results showing that culling vampire bats (a host species but not virus specific intervention) reduced BIV transmission, contrasts with the observed effects of the same culling program on rabies transmission, where culls were inconsequential for spillovers to livestock and accelerated the spatial spread of rabies when applied in areas with active viral circulation (32,62,63). We speculate that differing outcomes of the same intervention on co-circulating viruses reflects differences in viral maintenance strategies, with rabies transmission influenced more by host dispersal than density and BIV transmission being density dependent (11). From an applied perspective, our results provide the first evidence that non-specific interventions have unintended consequences on non-target viruses. It will be important for such interventions to be implemented with sufficient understanding of viral diversity to anticipate potentially counterproductive outcomes, particularly where multiple potential zoonoses co-circulate. Long term sample banks, growing capacity for multi-pathogen diagnostics (e.g., metagenomics and multiplexed serology), and the modelling framework we provide here may facilitate investigations into the dynamics of multiple pathogens.

As expected, several areas of uncertainty accompany these results. First, the data were sparse and inconsistent at the bat roost-level over both time and space, necessitating that we examine dynamics at the larger spatial scale of zones. Even so, the North zone in particular had significant stretches of time in which no data were available (> 3 years). However, temporal sampling biases are unlikely to produce the observed differences in seasonality between zones as sampling took place throughout the year, with at least some samples collected in all zones each month except January. Further, both feedback between the individual-level data and the model mechanism, and the use of data from all three zones for the key within-host parameters makes these estimates robust and provide a flexible framework for the irregular data common to wildlife studies. Another limitation is that our models were unable to include stochasticity (leading to the post-hoc exploration of stochastic viral extinction), direct links between zones to explain multi-annual synchrony, or an external force of infection due to computational power issues. Finally, BIV genomes were not detected in our limited metagenomic sequencing, so we cannot confirm whether the antibody responses that we detected are in response to the same H18 previously detected in Peruvian bats, a still undiscovered subtype, or a combination (64). Thus, ‘reinfections’ in model 1 could be as a result of reinfection with cross-reactive but different strain rather that the originally infecting strain.

In summary, by generating a Bayesian framework to incorporate irregularly collected field data from samples into mechanistic models, we provide the first projection of BIV transmission dynamics in a putative bat reservoir. Our results point to vampire bats as a competent reservoir of influenza, which may operate independently of or in conjunction with more widely cited fruit bat hosts, highlighting the need for *in vitro* and *in vivo* studies of this geographically widespread and ecologically connected bat. We further demonstrate the value of analysing anthropogenic interventions to verify presumptive reservoirs and establishing that non-pathogen-specific interventions can have unexpected consequences on off-target pathogens.

## Methods

### Data collection

The transmission dynamics of BIV were investigated based on longitudinal data collected from 30 bat roosts within 6 administrative Regions of Peru, between 2007 and 2018. Sampling in each bat roost was irregular over the study period. To obtain sufficient and relatively consistent sample sizes for model fitting, samples were grouped into three zones of Peru: North (combining Amazonas and Cajamarca colonies; N=7), Central (combining all colonies within Lima; N=4), and South (combining all bat colonies within Apurimac, Ayacucho, and Cusco; N=19; Figure 1).

Bats were captured using mist nets, harp traps or butterfly nets, then placed in individual cloth bags before processing. For serological assays, a maximum of 250μl of whole blood was collected by lancing the propatagial vein with a sterile 23-gauge needle. Blood was collected with heparinized capillary tubes, centrifuged in the field using serum separator tubes and stored on cold packs until it could be frozen (typically 0–3 days). Faecal samples for metagenomic sequencing were collected as rectal swabs, preserved in RNAlater, as described in Bergner et al (31).

Capture and sampling of bats was approved by the Research Ethics Committee of the University of Glasgow School of Medical, Veterinary and Life Sciences (Ref081/15) and by the University of Georgia Animal Care and Use Committee (A2014 04-016-Y3-A5). Field collections were authorized (per the efforts of the NGO Illariy) by the Peruvian government (103-2008-INRENA-IFFS-DCB, RD-222-2009-AG-DGFFS-DGEFFS, RD-299-2010-AG-DGFFS-DGEFFS, RD-009-2015-SERFOR-DGGSPFFS, RD-264-2015-SERFOR-DGGSPFFS, RD-142-2015-SERFOR-DGGSPFFS, RD-054-2016-SERFOR-DGGSPFFS).

### Detection of BIV antibodies

Indirect ELISA assays were performed by adapting the protocol established by Tong et al. (17) as follows to detect the presence of BIV IgG antibodies in bat serum. No commercial controls were previously tested against bat serum; therefore, the assay was initially developed using recombinant hemagglutinin H18 protein (rHA 18) and positive sera controls supplied by the CDC. From these initial tests, the recombinant Influenza A H18N11 (A/flat-faced bat/Peru/033/2010) Hemagglutinin / HA Protein generated in baculovirus (catalogue number 40324-V08B, Sino Biologicals) gave the best response to previously tested positive bat serum, and was used to coat the ELISA plate in subsequent ELISA assays at the same concentration of 1 μg/mL (100 μL per well) in PBS, pH 7.4 overnight at 4°C. Baseline negative controls were produced by pooling bat sera obtained from sacrificed bats that were non-reactive to rHA 18 into single assay tubes which were stored at −20°C for single use with each ELISA plate. For later assays, a commercial polyclonal antibody generated in rabbit (Influenza H18N11 (A/flat-faced bat/Peru/033/2010) Hemagglutinin / HA1 Antibody, Rabbit PAb, Antigen Affinity Purified, catalogue number 40324-T62, Sino Biologicals) was included as a positive serum control, diluted at 1:1000 in 2.5% non-fat milk-PBS with Tween 20 (0.1%), pH 7.4. The cut-off point of detectable response (based on 490 nm at 0.1 s OD) for the ELISA was modified to values above the average negative control plus 2 standard deviations. This generated a time series of seropositive and seronegative bats per sampled month for each zone (Figure 1).

### Metagenomic sequencing

The metagenomic sequences used in this study were previously published by Bergner et al. (2020) and are available in the European Nucleotide Archive (Project number PRJEB34487) (31). Faecal samples from 172 bats were collected between 2013-2017 (Figure 1) from 11 bat colonies within the three zones used in this study. RNA extracted from these samples was pooled by the sample collection site (1 pool of 5; 1 pool of 7; 16 pools of 10) and sequenced using the Illumina NextSeq500. The resulting reads were processed through a bioinformatics pipeline and compared to the Viral RefSeq Protein NCBI database. Additionally in this study, reads from each pool were aligned to reference sequences for each segment of an H18N11 genome published by Tong et al. (CY125949) using bowtie2 (17,65).

### BIV transmission dynamics

We fitted Bayesian compartmental models to the longitudinal BIV seroprevalence and metagenomic data collected from *D. rotundus* bats from 3 zones of Peru between 2007-2018. Our base compartmental model (“waning immunity”) followed an SEIRS structure, as used to model IAV transmission in other species, including birds and humans (66–68). This SEIRS model assumed that all bats transition from susceptible (S) to exposed (E) upon infectious contact with an infected bat at rate β, then progress to an infectious state after δ days (I). Bats then recover (R) with immunity (and thus are seropositive) after an infectious period of α days and remain immune for γ days after which they become susceptible once more (equations 1-4). Bats are born at rate *b* proportional to the total population size (N) and natural death occurs at rate *d* from each compartment. We assume here that the presence of antibodies is equal to immunity, and that antibody detections result from a single circulating strain of BIV, as opposed to multiple interacting strains.

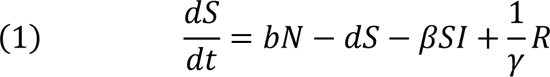

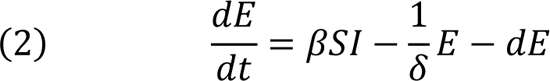

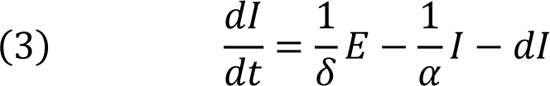

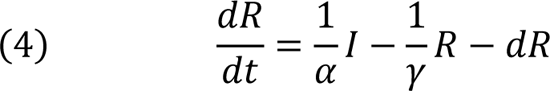

The “lifelong immunity” model followed an SEIR^+^R^-^ structure, where compartments and transitions S-E-I-are the same as in model 1. Upon recovery, individuals become seropositive (R+), and remain so for γ days, at which time antibodies wane below the detection threshold, but the bat remains immune (R-). Individuals can return to R+ due to immune reactivation upon meeting an infected bat at rate β, but do not return to the susceptible compartment.

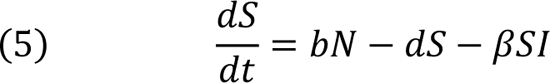

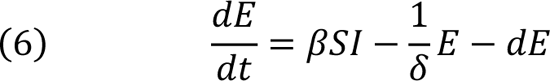

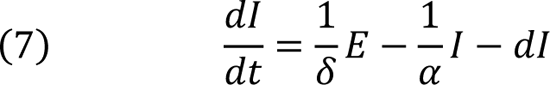

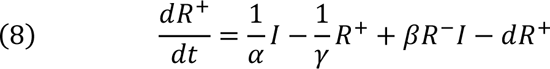

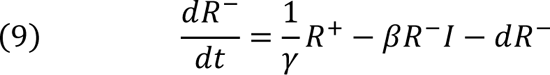

In the ‘South’ zone, an additional component was included in the model to evaluate the impact of bat culling that took place in this zone during the study period, from 2014-2016. In our model, bats were culled from all compartments at a rate (*csf*) proportional to the total number of bats recorded as being culled in each month *t* during the study period (*cull*[*t*]).

For example, for the infectious state I:

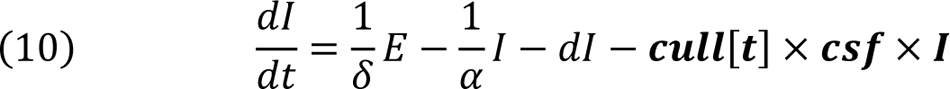

Additionally, we attempted to incorporate (a) environmental transmission and (b) transmission between zones; however, these models failed to converge, and posterior checks were poor.

The following assumptions were made on the population dynamics of *D. rotundus*. We assumed that all bats were born susceptible to BIV, without protection from maternal antibodies. Without the influence of outside factors, we also assumed that the birth rate was equal to the natural death rate, such that the population size of each zone remained constant throughout the study period. We did not have estimates of colony size for the majority of sampled colonies, nor of the total number of unsampled colonies with which to estimate the population of the zones. As such, we used a deterministic framework for this model fitting, with a total population of 1. We modelled transmission as density dependent, as is usual for influenza viruses in other species (66).

Most parameters in our models were given informative priors based on published estimates following experiments on captive animals (Table 1). The exceptions were the transmission rate β and the immune period γ. To understand the short-(seasonality) and long-term periodicity of BIV, β was constructed through a combination of two sine waves, with periodicity confined to this term in order to avoid presumptuously assigning causation (e.g., instead of including a periodic birth function). All periodic parameters within β(*t*) were estimated separately for each zone, apart from period_1_ which was fixed at 365 days to evaluate the influence of seasonal forcing, although the timing (*x*_1_) and magnitude (β_1_) of this period were estimated.

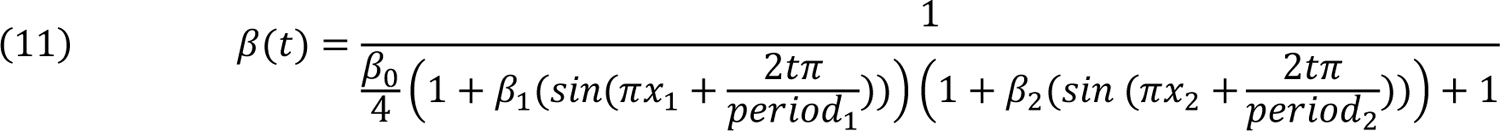

### Occupancy-model for estimation of immune period **γ**

To inform the immune period γ we used the individual-level longitudinal data, collected from individuals recaptured multiple times over the study period. Given the starting serostatus (0 or 1) of a resampled bat *i* at sample point *T*1(*B*_i,T1_), a simple occupancy model predicted the probability of this individual being seropositive at sample point *T*2 based on the value of γ (antibody waning; an individual transitions from 1->0), and the probability of infection *pc*_t_ based on the total proportion of bats that became Infected in the intervening time from the trajectories of the ODEs, (infection; 0->1):

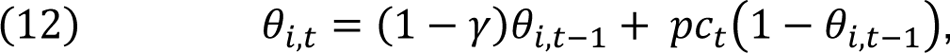

where θ_i,t_ is the probability of an individual being seropositive and is updated for each time point between sampling points *T*1 and *T*2. The likelihood of seropositivity at the second sampling point for each individual was defined through a Bernoulli distribution:

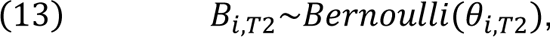

where *B*_i,T2_ is the serostatus (0 or 1) of recaptured bat *i* at its second sample point T2, and θ_i,T2_ is the probability of a bat being seropositive at this point. For bats that were sampled 3 times or more, consecutive samples were divided into pairs such that a bat sampled 3 times has samples pairs (*T*1, *T*2) and (*T*2, *T*3). These observations were assumed to be independent.

### Model fitting to population-level data

The proportion of seropositive bats was predicted for each zone by solving the model on a daily timescale. The likelihood of the model described the proportion of seropositive bats averaged for each 30-day period in a zone MR, through a binomial likelihood function:

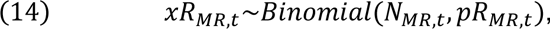

where *x*R_MR,t_ is the number of seropositive bats from zone MR at sample time t among total sample size N_MR,t_ and pR_MR,t_ is the probability of being seropositive (R) estimated by the model.

A limited number of faecal samples previously tested by metagenomic sequencing were also included in the model fitting (31). These samples were used to evaluate the likelihood of sampling active infection across the entire sampling period where the number of positive bats followed a binomial distribution:

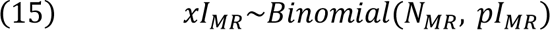

Where *x*I_MR_ is the total number of sequence-positive bats from each zone during the sampling period among total sample size N_MR_, and pI_MR_ is the probability of being infected (I) for a given zone estimated by the model.

### Model evaluation and selection

All models were fitted using the software JAGS (69) interfaced with R version 4.2.3 (70) using the package rjags (71). The models assuming waning and lifelong immunity were run with 4 chains in parallel for 100,000 iterations with a burn-in of 50,000 and a thin interval of 50 for a total of 1000 samples per chain for memory saving purposes, then compared based on the deviance information criterion (DIC) and leave-one-out information criterion (LOOIC (72)), convergence, and model fit (Table S2). Models comparing data inclusion were run with 4 chains in parallel for 5000 iterations with a burn-in of 2500 and a thin interval of 2 such that there were a total of 1250 samples. We inspected the models for chain convergence by visualisation of trace plots, Gelman-Rubin convergence diagnostic (<1.1 considered to demonstrate convergence), and effective sample size. To assess model fit we compared the observed number of seropositive bats to the posterior distribution of the expected number of seropositives estimated from each of the MCMC samples. For each time point at which data was available, we calculated the percentile where the data point fell within the cumulative distribution function (Figure 1 and Figure S3).

### Sensitivity analysis via simulations

A sensitivity analysis was carried out to test (a) the accuracy of model fitting over a range of parameter values, and (b) the impact of including different data types on this model fitting, using 96 randomly generated parameter sets. The population-level test data was generated by running a deterministic simulation of the model (using the deSolve package in R (73)), then using the true sample sizes from the data at each time point at which collection occurred to produce the expected number of seropositive bats. This was repeated for all three zones of Peru. Longitudinal and metagenomic datasets were also simulated using the output of the deterministic model. Model fitting was then carried out using JAGS with either the population-level serology simulated data only, or with all three simulated datasets. For each set of parameters, we recorded the mean and standard deviation output from the posterior distribution of the jointly estimated parameters, and of the regionally estimated parameters for the South zone. The final model (waning immunity) was run with and without the individual-level serology and metagenomic data in order to evaluate its influence on parameter estimates.

### Stochastic simulations

To identify the vampire bat population size required to maintain BIV independently, a series of stochastic simulations were run using the adaptivetau package in R (74) using the mean transmission parameters taken from the posterior distribution of the model fitting. For each zone, 250 simulations of the stochastic models were run for total population sizes ranging from 2500 to 150,000 bats, and the proportion of simulations in which BIV was maintained (at least one individual in the exposed or infected class) until the end of the 12-year simulated study period was recorded.

## Supporting information

Full Supplementary materials

## Acknowledgements

This work was funded by the Wellcome Trust (Sir Henry Dale Fellowship: 102507/Z/13/A; Senior Research Fellowship: 217221/ Z/19/Z), the US National Science Foundation (DEB-1020966; DEB-2011069), the BBSRC (BB/V003798/1) and the Leverhulme Trust (RPG-2015259). MV was supported by European Research Council under the European Union’s Horizon 2020 Research and Innovation Programme (grant 852957). We are also grateful to the field assistants who carried out bat capture and sampling in Peru.

## Data availability

Sample data, as well as all code used to fit each transmission model, analyse results, and plot figures, will be available on GitHub upon publication.

## Notes

### Competing Interest Statement

The authors have declared no competing interest.

