## Supplementary material for "Dynamics of influenza transmission in vampire bats revealed by longitudinal monitoring and a large-scale anthropogenic perturbation": Full Supplementary materials

^5^Asociación para el Desarrollo y Conservación de los Recursos Naturales (Illariy)

^6^Departamento de Mastozoología, Museo de Historia Natural, Universidad Nacional Mayor de San Marcos, Lima, Perú

^7^Centro de Investigaciones Tecnologicas, Biomedica y Medioambientales-CITBM, Faculty of Medicine San Fernando, Universidad Nacional Mayor de San Marcos, Lima, Perú

† Joint senior authors

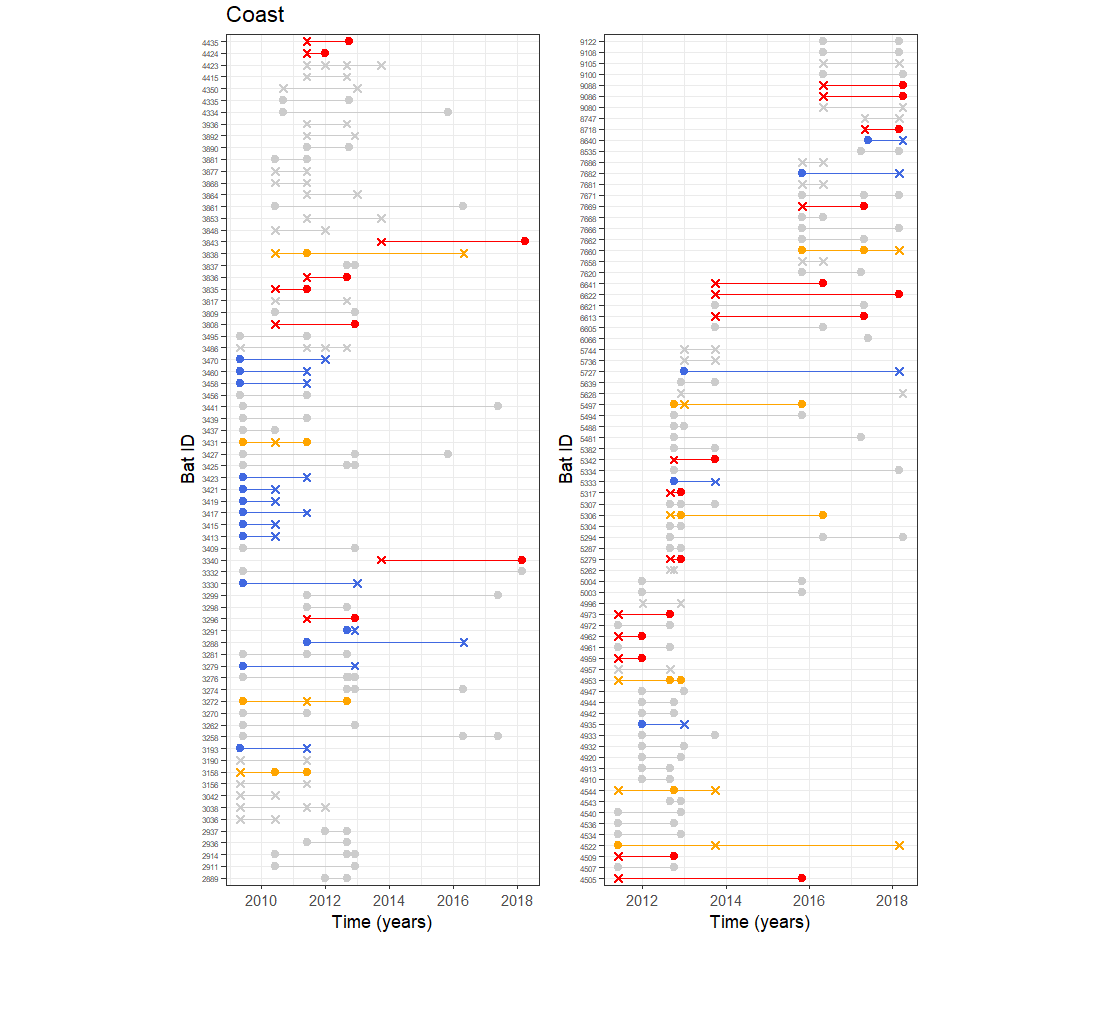

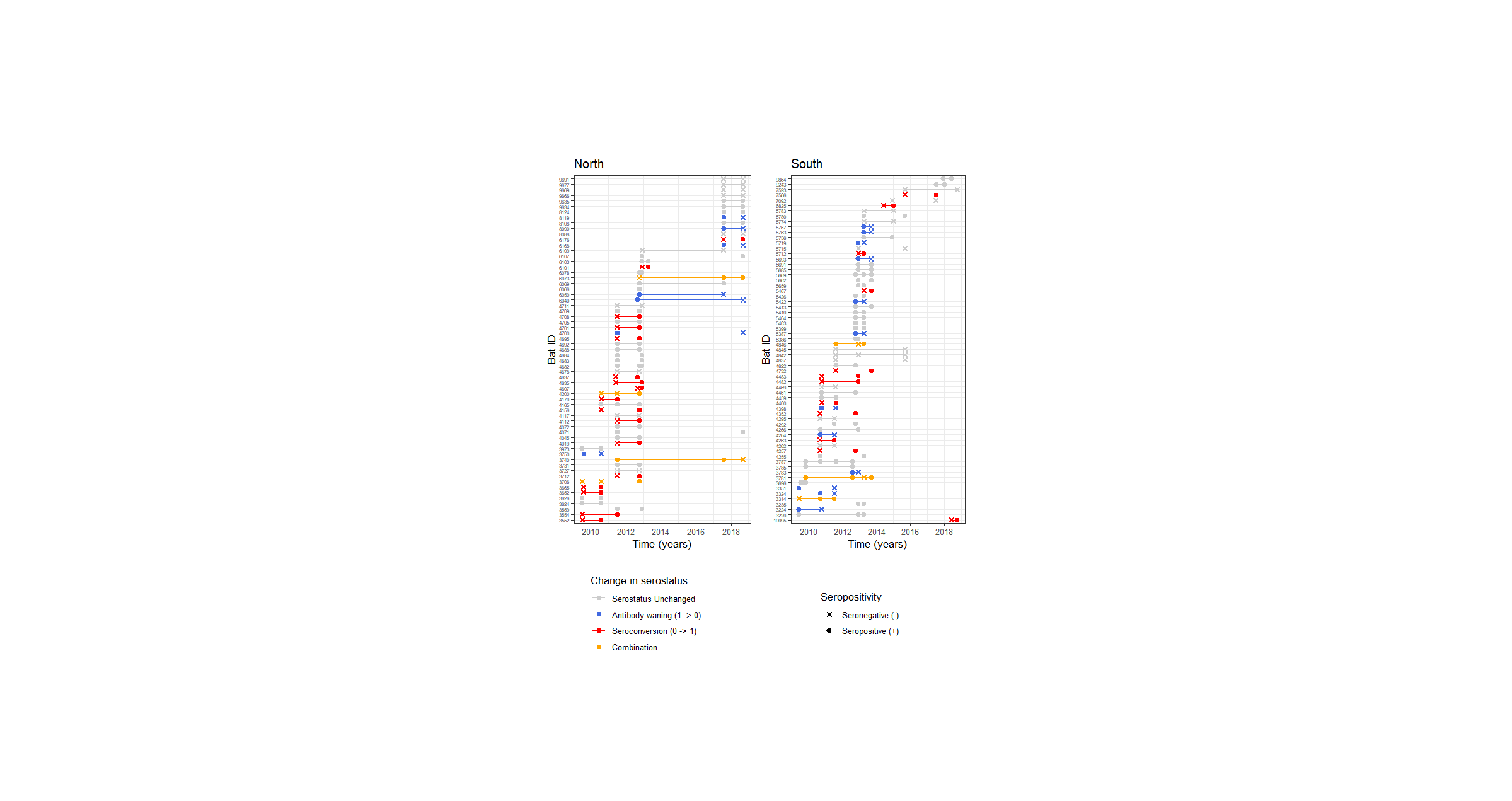

Figure S1 – Longitudinal serology data from recaptured bats for each zone (Central N=149; North N=63; South N=65).

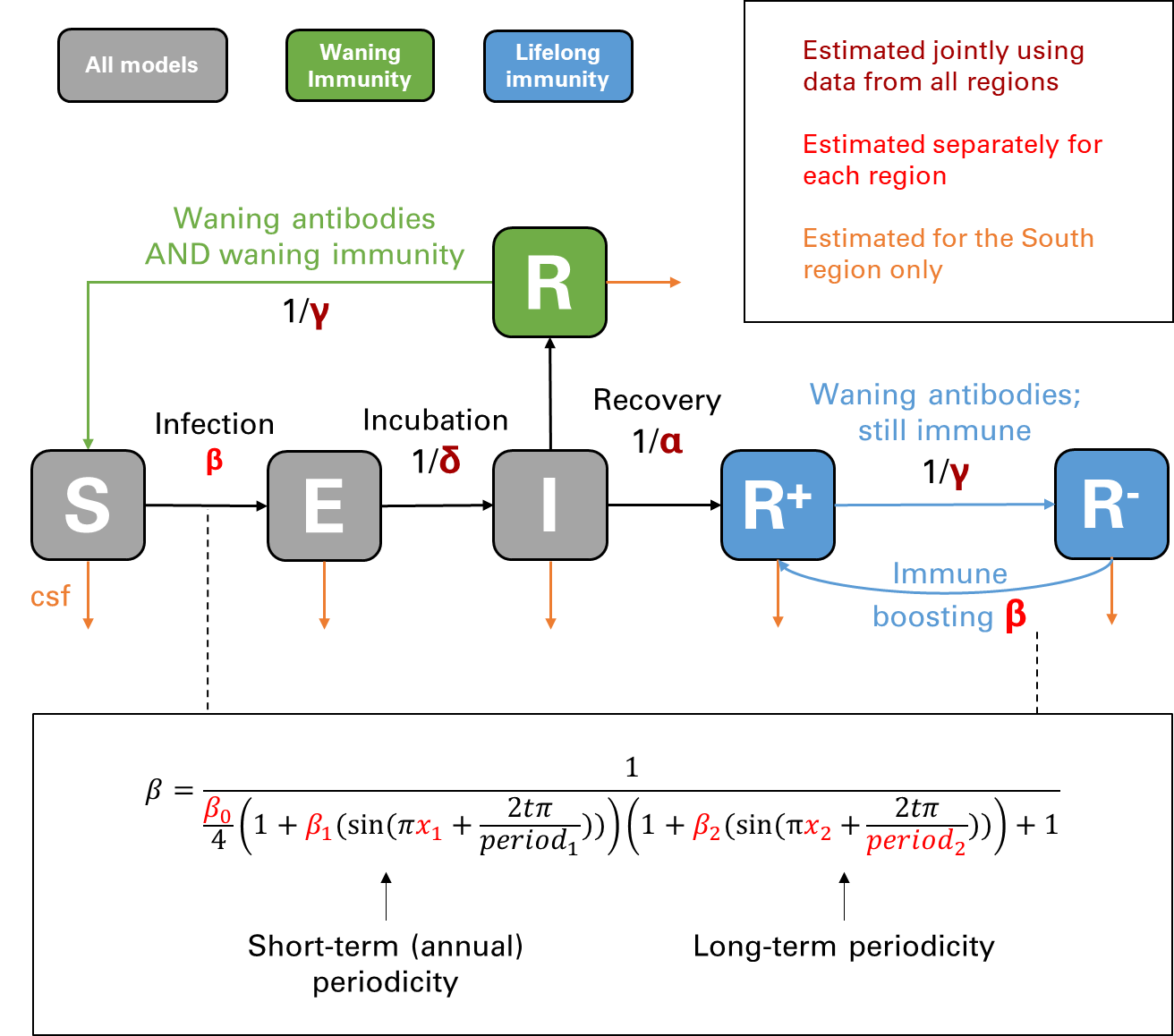

Figure S2 – Schematic of BIV compartmental models for both waning and lifelong immunity. Grey compartments and black arrows show the states/rates of transitions that are common to all models/zones (S=Susceptible, E=Exposed, I=Infected). Green shows the states and transitions for the model assuming waning immunity where R=Recovered and seropositive. Blue shows the states and transitions for the model assuming lifelong immunity where R^+^=Recovered and seropositive and R^-^=Recovered but seronegative.

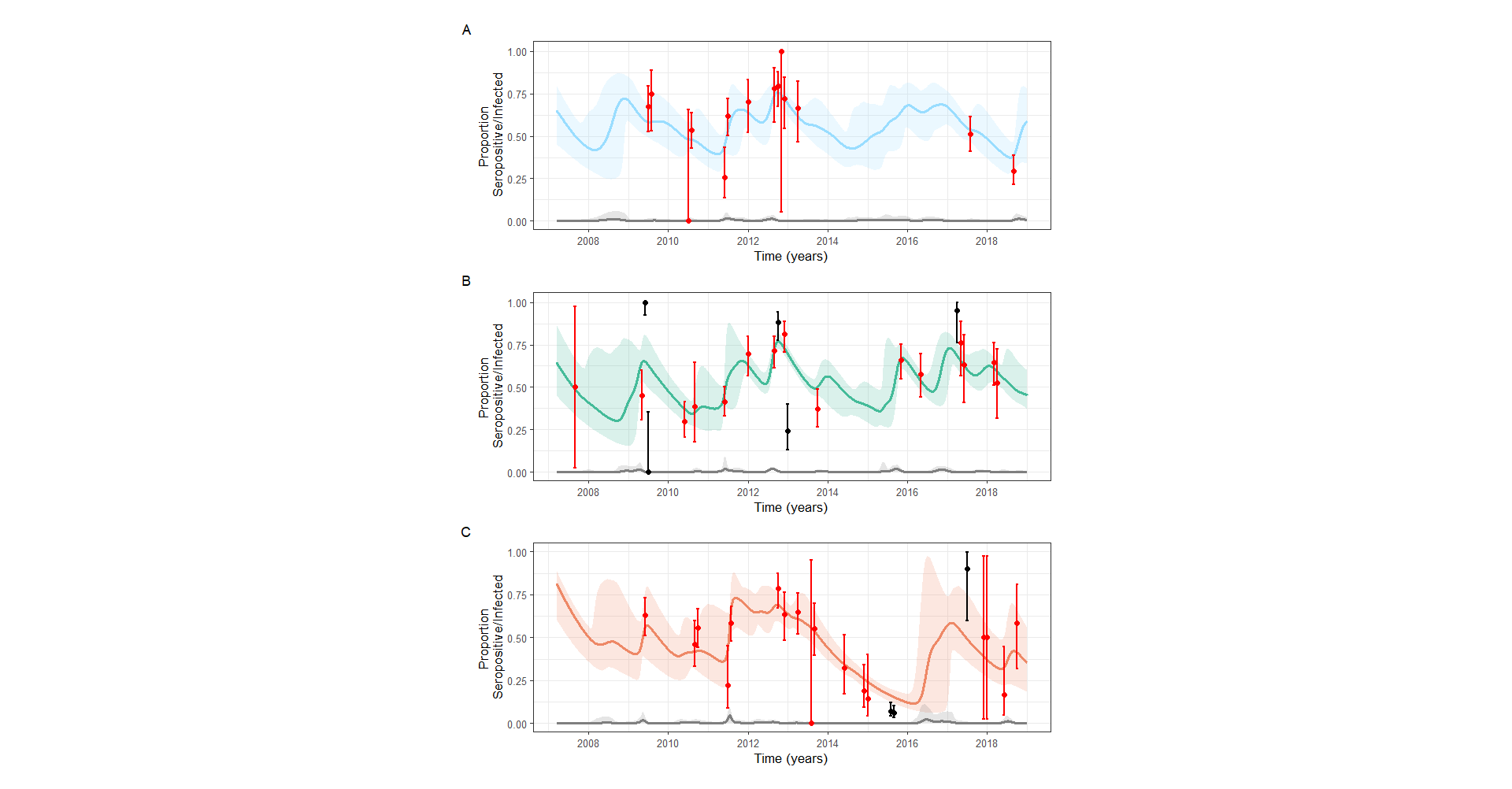

Figure S3 – Bat Influenza virus seroprevalence data and posterior predictive distributions when assuming lifelong rather than waning immunity. Population-level seroprevalence data with binomial confidence intervals per month for populations in (A) North, (B) Central, and (C) South are shown with colours indicating whether the observed seropositivity falls inside (red) or outside (black) the 95% CI of the posterior cumulative distribution function. The mean trajectory for the Recovereds fit by the model is shown for each population by the coloured solid line, and the 95% CIs by the shaded regions. The mean trajectory for the Infecteds fit by the model is shown for each population by the grey line and shaded region.

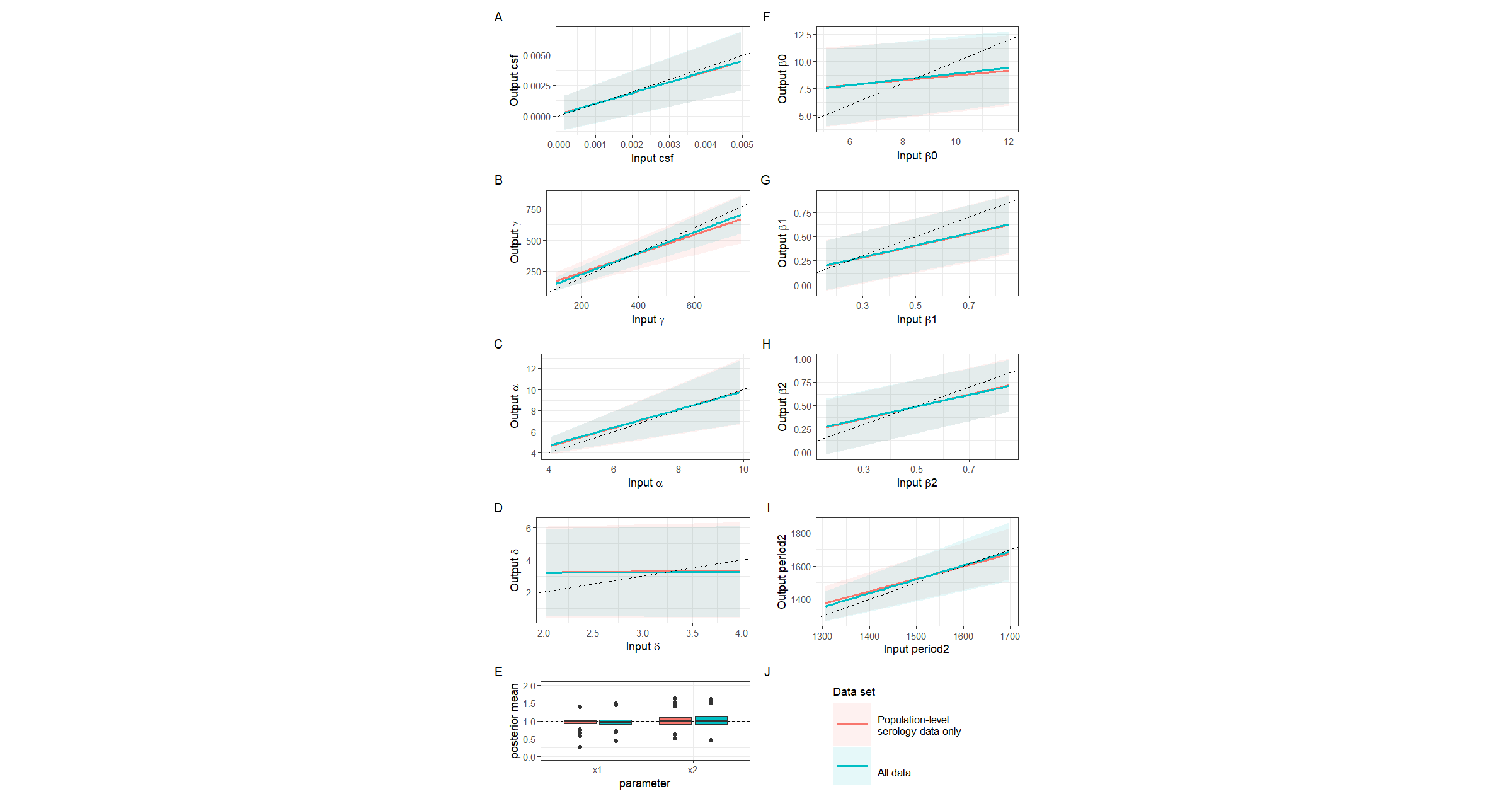

Figure S4 – Model fitting to test data and data inclusion comparison. The x-axis of each graph shows the input parameter value, and the y-axis shows the mean of the posterior distribution for that parameter when model fitting to test data. The shaded areas show 95% credible intervals. The dotted line on each plot is at x=y where the model prediction matches the input value of the estimated parameter. Mean lines which are closer to this dotted line show better prediction by the model.

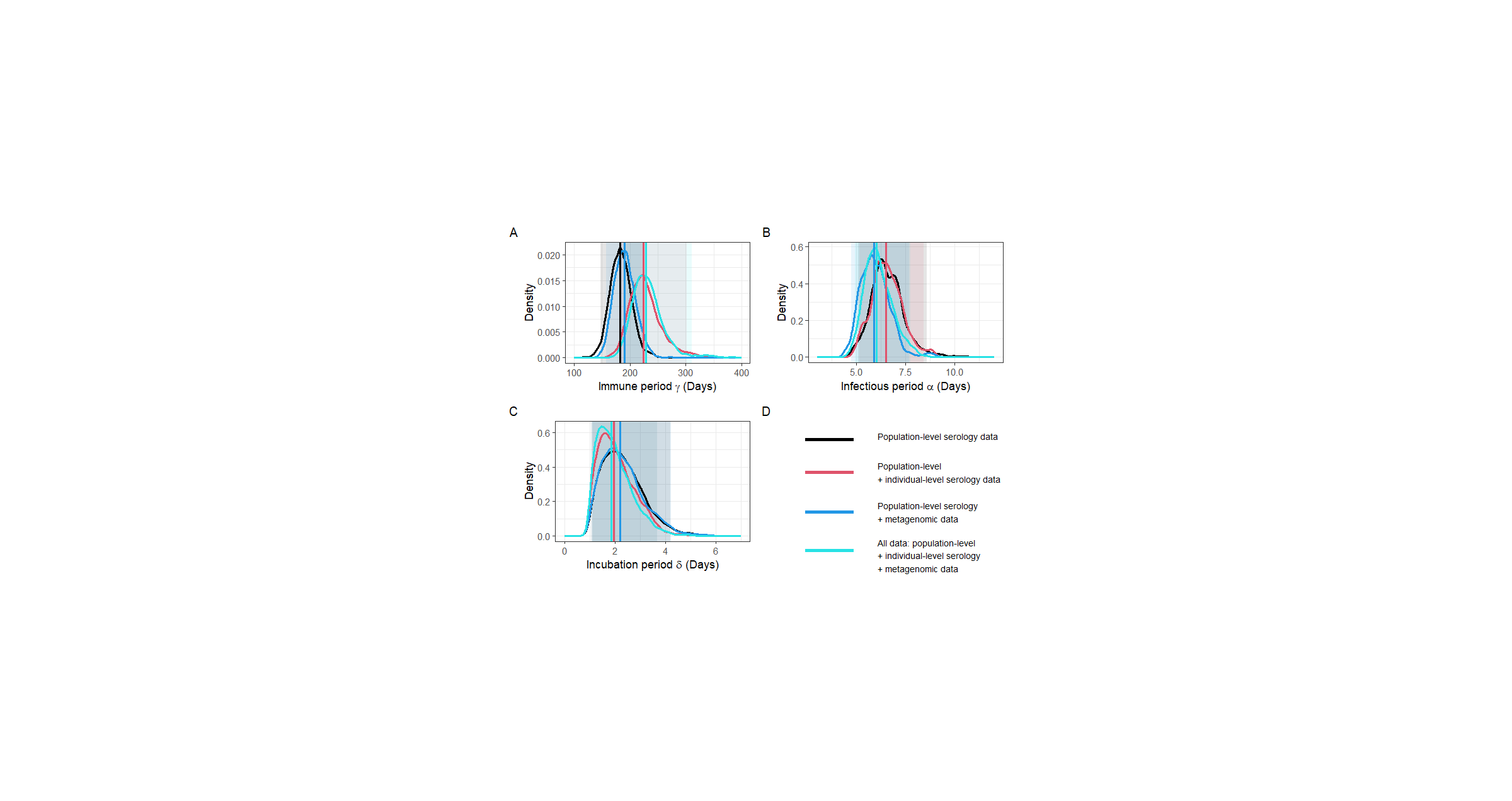

Figure S5 – A comparison of the posterior distributions of jointly estimated parameters when different data types are included in the model. Plots show the posterior mean and distribution when using different combinations of data, including population-level serology, individual-level seroconversion histories, and metagenomic data. Vertical lines show the mean of the distribution, and shaded areas show the 95% credible intervals.

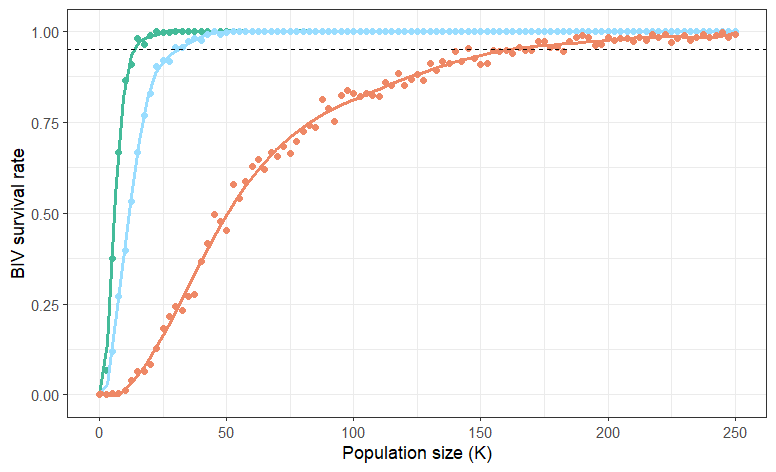

Figure S6 – Population size required for long term maintenance of BIV in vampire bat populations. Points show the proportion of stochastic simulations (out of 250 simulations) in which BIV survives for the full sampling period in each zone (green=Central; blue=North; orange=South). The dotted line shows the population size needed for 95% of simulations to result in BIV survival, where at least one Infected or Exposed bat is present in the population at the end of the 12-year simulation. For the Central zone, this is ~15,000 bats, for the North ~30,000 bats, and for the South ~155,000 bats. The larger population size requirement in the South reflects the impact of culling.

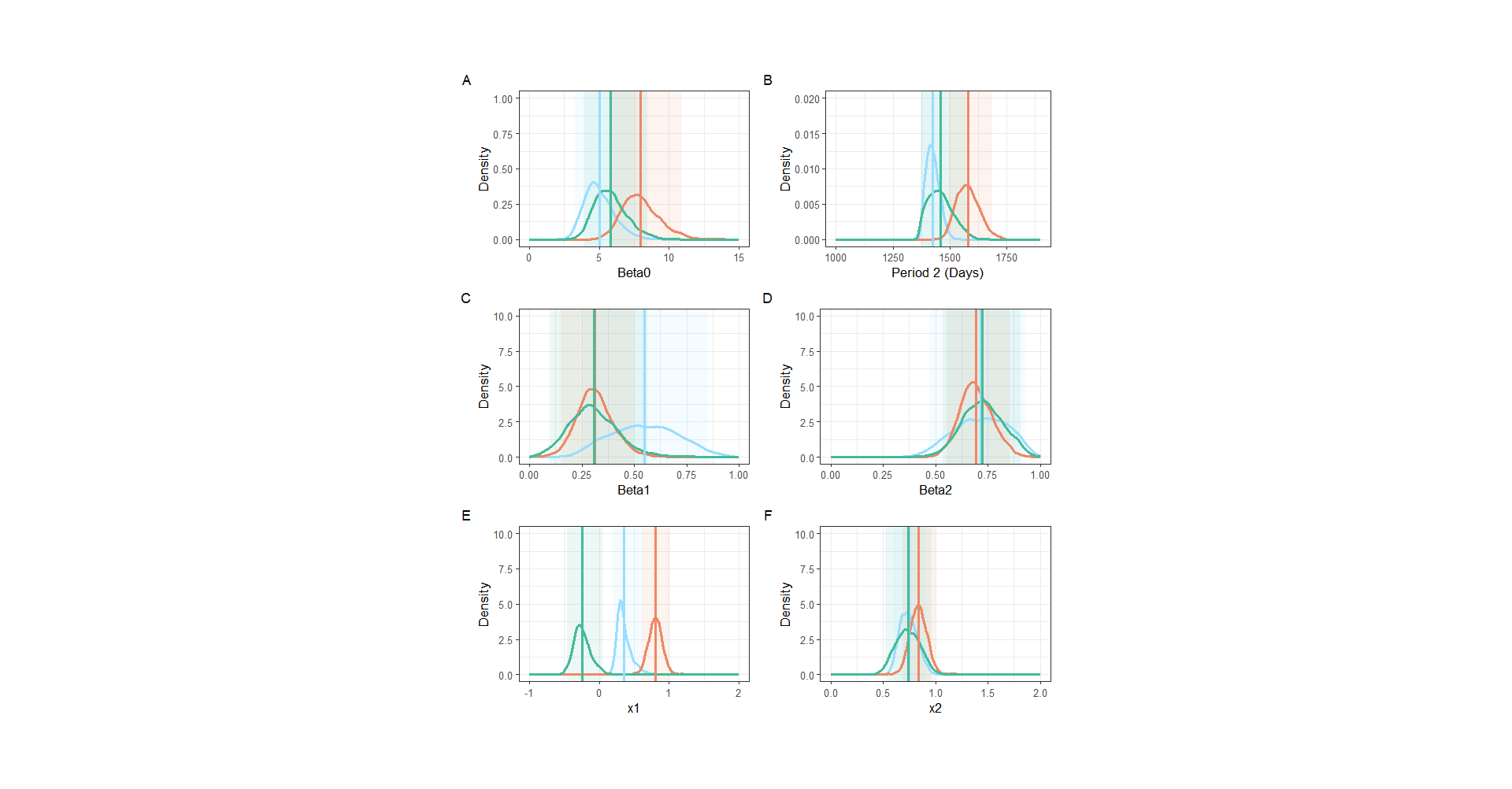

Figure S7 – Posterior distributions for all parameters within the formula for the rate of transmission β (Equation 11). β = base rate of transmission, beta1 = magnitude of short-term cycle peaks, beta2 = magnitude of long-term cycle peaks, period2 = the time between peaks of the long-term cycles, x1 = timing of the short-term cycles, and x2 = timing of the long-term cycles. Posterior means for each zone (blue=North; green=Central; orange=South) are shown by the vertical lines, and 95% credible intervals are shown by the shaded areas.

Table S1 – Details of all sites from which bats samples used in this study were taken. DR = *Desmodus rotundus*. Direct culling refers to sample sites within areas where Vampiricide was applied to captured bats.

| Site abbreviation | Region | Zone | Years sampled | Total DR sampled | Total DR positive | Sero-prevalence | Other bat species | Direct culling |
| --- | --- | --- | --- | --- | --- | --- | --- | --- |
| AMA1 | Amazonas | North | 2011 | 13 | 10 | 0.77 | Y | N |
| AMA2 | Amazonas | North | 2011-2013 | 36 | 21 | 0.58 | Y | N |
| AMA3 | Amazonas | North | 2011 | 3 | 2 | 0.67 | N | N |
| API1 | Apurimac | South | 2009-2015, 2017-2018 | 227 | 119 | 0.52 | Y | Y |
| API13 | Apurimac | South | 2009-2015 | 115 | 71 | 0.62 | Y | Y |
| API140 | Apurimac | South | 2010-2015, 2017 | 161 | 59 | 0.37 | N | N |
| API141 | Apurimac | South | 2015 | 45 | 5 | 0.11 | N | Y |
| API15 | Apurimac | South | 2015 | 7 | 0 | 0 | N | N |
| API16 | Apurimac | South | 2015 | 14 | 0 | 0 | N | Y |
| API17 | Apurimac | South | 2015 | 31 | 1 | 0.03 | N | Y |
| API18 | Apurimac | South | 2015 | 9 | 2 | 0.22 | N | Y |
| API3 | Apurimac | South | 2010-2013 | 70 | 39 | 0.56 | N | Y |
| API9 | Apurimac | South | 2009-2013 | 80 | 46 | 0.58 | N | Y |
| AYA1 | Ayacucho | South | 2015 | 15 | 0 | 0 | N | Y |
| AYA11 | Ayacucho | South | 2015 | 23 | 5 | 0.22 | Y | N |
| AYA13 | Ayacucho | South | 2015 | 6 | 1 | 0.17 | N | Y |
| AYA14 | Ayacucho | South | 2015 | 32 | 4 | 0.13 | N | Y |
| AYA15 | Ayacucho | South | 2015 | 29 | 1 | 0.03 | Y | Y |
| AYA4 | Ayacucho | South | 2015 | 53 | 1 | 0.02 | N | Y |
| AYA7 | Ayacucho | South | 2015 | 6 | 0 | 0 | Y | Y |
| CAJ1 | Cajamarca | North | 2009-2012, 2017-2018 | 117 | 64 | 0.55 | Y | N |
| CAJ2 | Cajamarca | North | 2009-2012, 2017-2018 | 181 | 115 | 0.64 | Y | N |
| CAJ3 | Cajamarca | North | 2009-2012, 2017-2018 | 118 | 59 | 0.50 | Y | N |
| CAJ4 | Cajamarca | North | 2010-2013, 2017-2018 | 146 | 72 | 0.49 | Y | N |
| CUS6 | Cusco | South | 2015 | 31 | 0 | 0 | Y | N |
| CUS8 | Cusco | South | 2015 | 25 | 1 | 0.04 | N | N |
| LMA10 | Lima | Central | 2007, 2009-2013, 2015 | 130 | 30 | 0.23 | N | N |
| LMA4 | Lima | Central | 2009-2013, 2015, 2017-2018 | 171 | 125 | 0.73 | Y | N |
| LMA5 | Lima | Central | 2009-2013, 2015-2018 | 221 | 118 | 0.53 | N | N |
| LMA6 | Lima | Central | 2009-2013, 2015-2018 | 436 | 296 | 0.68 | Y | N |

Table S2 - Comparison and evaluation of the models assuming waning and lifelong immunity.

|  | **Waning Immunity** | **Lifelong immunity** |
| --- | --- | --- |
| Structure | SEIR | SEIR^+^R^-^ |
| DIC | 836.90 | 923.22 |
| looic estimate | 867.30 | 1013.40 |
| looic se | 85.60 | 101.40 |
| elpd-loo estimate | -433.70 | -506.70 |
| elpd-loo se | 42.80 | 50.70 |
| Max Rhat | 1.02 | 2.76 |
